# Mut-BPE: A Modified BPE Strategy Improves Variant Effect Prediction

**DOI:** 10.1101/2025.12.01.691503

**Authors:** Yucheng Xu, Weicai Long, Yawen Lu, Xu Yang, Yanlin Zhang

## Abstract

Byte Pair Encoding (BPE) is widely used in genome foundation models for its ability to compress long DNA sequences into fewer tokens. However, its variable-length tokens often span multiple nucleotides, limiting the model’s sensitivity to single-nucleotide variations—an essential requirement for accurate Variant Effect Prediction (VEP). We introduce Mut-BPE, a training-free, plug-and-play tokenization strategy that augments BPE with explicit single-nucleotide resolution at variant sites. Mut-BPE preserves the efficiency of BPE while enhancing its ability to represent subtle genomic alterations. To evaluate its effectiveness, we applied Mut-BPE to DNABERT-2 and conducted extensive experiments across six diverse datasets spanning gene expression, pathogenicity, and trait-associated variants, under zero-shot and fine-tuning settings. Mut-BPE consistently outperformed conventional BPE tokenization, yielding significant improvements in both AUROC and AUPRC, particularly in imbalanced datasets. These results highlight Mut-BPE as a practical enhancement for genomic foundation models in VEP tasks. Code availability: https://anonymous.4open.science/r/Mut-BPE-1182

## I. Introduction

Accurately predicting the functional impact of genomic variants remains a core challenge in precision medicine [3], drug discovery [1], and genetic disease diagnostics [2]. Among all variant types, single nucleotide polymorphisms (SNPs) and insertions/deletions (Indels) are major contributors to inherited disorders, inter-individual variation in drug response, and evolutionary processes [4]. Traditional variant effect prediction (VEP) methods, such as CADD [6] and phyloP [7], have provided valuable insights by leveraging functional annotations or evolutionary conservation. However, these approaches often fall short on large-scale genomic datasets due to their limited ability to capture complex, long-range dependencies in DNA sequences [8].

The emergence of deep learning, particularly transformer-based models, has significantly advanced DNA sequence modeling. Pre-trained genome foundation models can learn rich, context-aware representations from large-scale unlabeled DNA sequences and be fine-tuned for downstream tasks, including VEP [9], [10]. A critical component of such models is the tokenization strategy used to convert raw DNA into model-readable inputs. Among various strategies, Byte Pair Encoding (BPE) has gained widespread adoption in models like Gena-LM [12], GROVER [13], and DNABERT-2 [11], due to its ability to compress sequences by merging frequent sub-sequences into variable-length tokens. This reduces input length and enhances computational efficiency, enabling the modeling of longer genomic contexts [14], [16].

Unlike fixed-length k-mer or character-level tokenizers, Byte Pair Encoding (BPE) produces variable-length tokens, where a single token can span multiple nucleotides. This design complicates the model’s ability to localize and represent single-nucleotide mutations at the token level. As a result, although BPE-based models excel at capturing broader sequence patterns, their lack of nucleotide-level resolution hinders their ability to accurately interpret subtle variations in DNA, such as single-base substitutions. This limitation is especially detrimental in VEP tasks, which require fine-grained discrimination of single-nucleotide variant effects—particularly in functionally critical regions like coding sequences or non-coding regulatory elements [15].

To address this limitation, we propose Mut-BPE, a simple, training-free, and plug-and-play modification to conventional BPE tokenization. Mut-BPE preserves BPE’s efficiency while introducing explicit single-nucleotide resolution at variant sites. Specifically, for each mutation, the token that spans the variant is split into three parts: the prefix, the single mutated nucleotide, and the suffix. This ensures that both the reference and alternative sequences share aligned token structures, differing only at the mutation site, thereby enabling more precise and mutation-aware representations.

We evaluate Mut-BPE by applying it to DNABERT-2 and conducting comprehensive experiments across six mutation datasets covering gene expression, pathogenicity, and trait-associated variants. Performance is assessed under zero-shot and fine-tuning settings. Our results show that Mut-BPE consistently outperforms standard BPE tokenization across all evaluation regimes, especially in imbalanced datasets. These findings underscore the importance of fine-grained tokenization in genomic modeling and position Mut-BPE as a practical and effective enhancement for BPE-based foundation models in variant effect prediction.

## II. Methods

### A. Datasets

To comprehensively evaluate the performance of DNA foundational models in Variant Effect Prediction (VEP) tasks, we utilized six datasets, as illustrated in Figure 1. These datasets span three primary categories: gene expression-related (eQTLs), pathogenicity-related, and complex trait-related mutations. For each variant within these datasets, we provide the reference DNA sequence, the single-base change of the mutated sequence, and its genomic coordinates (hg38 human reference genome). Crucially, each variant is associated with a specific label indicating its pathogenicity or functional impact. This diverse collection of data facilitates rigorous evaluation across zero-shot and fine-tuning experimental paradigms.

**Fig. 1.**
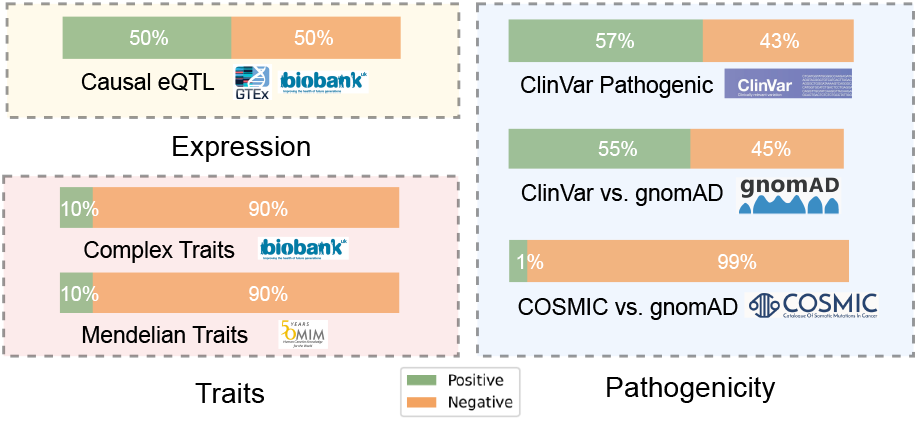
Introduction of VEP datasets: gene expression, pathogenicity and traits, and their labels distribution

Causal eQTL Dataset: Derived from the Enformer [17] model study and integrated into the Genomics Long-Range Benchmark [18], this dataset focuses on causal expression quantitative trait loci (eQTLs)—genetic variants directly influencing gene expression. Variants are labeled based on whether a single nucleotide polymorphism (SNP) perturbs gene expression, with binary labels assigned if the variant’s causal probability exceeds 0.9, as determined by population-based fine-mapping analysis. Chromosomes 9 and 10 serve as the test set, with other chromosomes forming the training set.

ClinVar Pathogenic Variant Dataset: This dataset leverages pathogenic mutations meticulously curated within the ClinVar database [20], a public archive of clinically relevant sequence variations and human phenotypes. Employed in the GPN-MSA study [19], this dataset involves classifying ClinVar pathogenic variants against ClinVar benign-labeled variants as controls. Chromosome 1 is designated as the test set in this study.

ClinVar Pathogenic vs. gnomAD Common Missense Variant Dataset: Also sourced from the GPN-MSA study and included in the Genomics Long-Range Benchmark, this dataset distinguishes ClinVar pathogenic variants from common missense variants found in the Genome Aggregation Database (gnomAD)—a vast collection of human genetic variation data [22]. The task involves classifying ClinVar pathogenic variants against common missense variants from gnomAD as controls. Chromosome 8 is utilized as the test set in this study.

COSMIC Frequent vs. gnomAD Common Missense Variant Dataset: Originating from the GPN-MSA work, this dataset focuses on somatic missense variants frequently observed across cancer tumors (from COSMIC [21], the Catalogue Of Somatic Mutations In Cancer) versus common missense variants from gnomAD. This dataset is characterized by a severe class imbalance, with a scarcity of positive instances, necessitating careful evaluation of metrics like precision and recall. Chromosome 1 is chosen as the test set in this study.

Mendelian Traits Dataset: Curated within TraitGym [23], this dataset represents one of two non-coding variant benchmarks for human genetics. It comprises causal variants for 113 Mendelian traits. For each putatively causal non-coding variant, nine non-coding control variants are sampled, meticulously matched by chromosome, consequence, and distance to the transcription start site (TSS). A minor allele frequency (MAF) less than 0.1% in gnomAD serves as a filtering criterion for pathogenic variants. Chromosome 11 is designated as the test set in this study.

Complex Traits Dataset: The second dataset from TraitGym consists of strong causal variant candidates across 83 complex traits. These putative causal variants are identified based on a posterior inclusion probability (PIP) greater than 0.9, derived from statistical fine-mapping results within UK BioBank data. Control variants are rigorously matched by chromosome, consequence, TSS distance, MAF, and linkage disequilibrium (LD) score in the UK BioBank. Chromosome 1 is used as the test set for this dataset in this study.

### B. BPE Tokenizer

Byte Pair Encoding (BPE) is a data-driven tokenization algorithm that iteratively merges the most frequent adjacent symbol pairs in a corpus to construct variable-length tokens. Originally developed for natural language processing, BPE has been widely adopted in genomic language models due to its ability to compress long DNA sequences and learn informative sub-sequence units [11], [12]. In the genomic context, BPE treats nucleotides (`A, C, G, T`) as individual characters and builds a vocabulary by repeatedly merging frequently co-occurring nucleotide pairs. The result is a flexible set of sub-word units that capture common DNA patterns while reducing the sequence length, which is beneficial for transformer-based models with quadratic attention complexity.

To illustrate how BPE operates, consider the following (Figure 2a). Given the reference sequence ATCAATGGCAATT, a trained BPE tokenizer may segment it into: [‘ATCA’, ‘ATGGC’, ‘AATT’], where each token represents a frequent sub-sequence learned from the pretraining corpus. If a single nucleotide within this sequence is substituted—for example, changing the guanine (G) at position 7 to adenine (A), resulting in the alternative sequence ATCAATAGCAATT—the same BPE tokenizer may produce a different segmentation: [‘ATCAA’, ‘TAGCAA’, ‘TT’]. This example demonstrates that even a single nucleotide change in the input sequence can result in a completely different set of BPE tokens, affecting both the content and number of tokens produced.

**Fig. 2.**
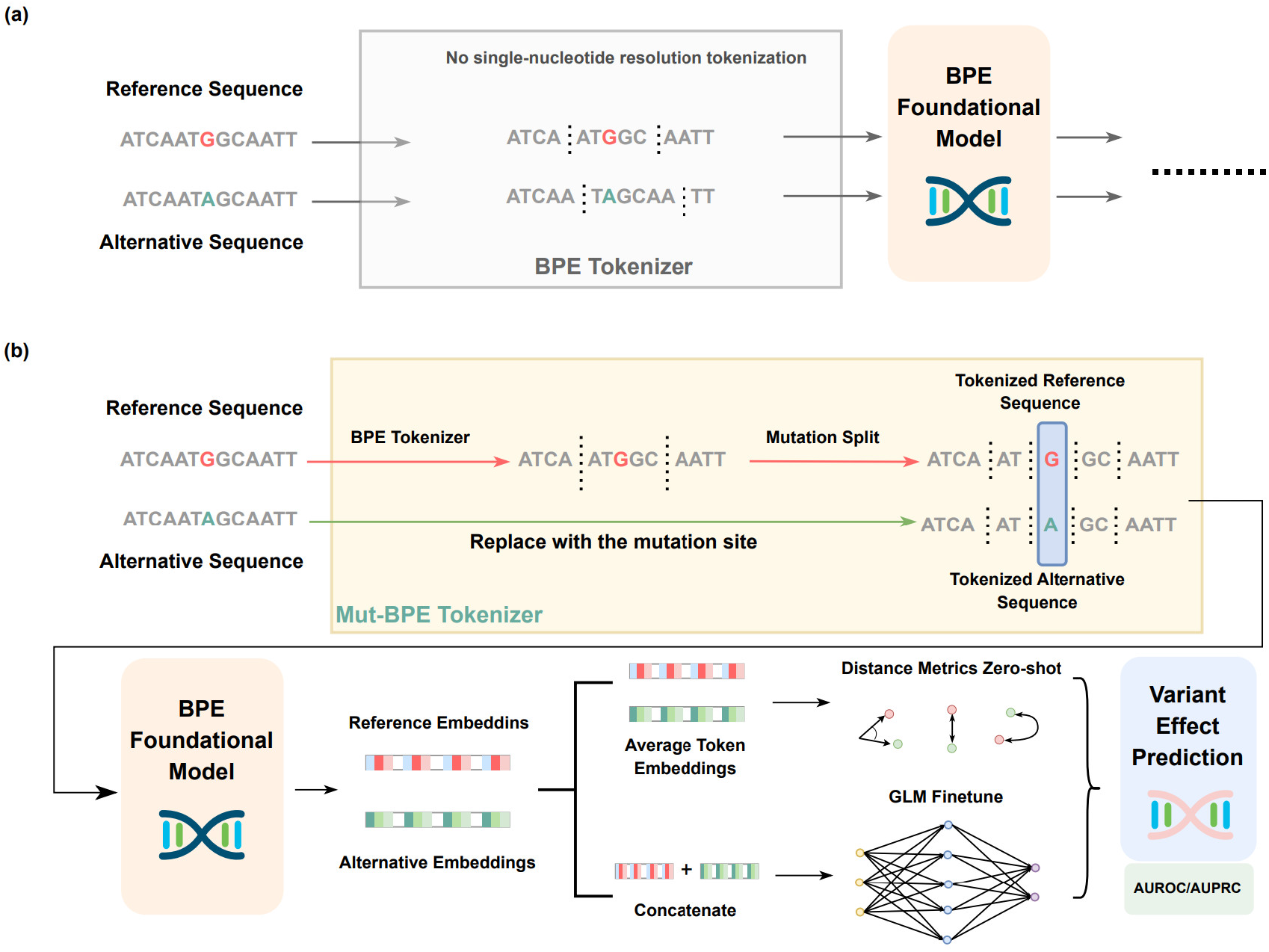
(a) is the daigram of single-nucleotide mutations tokenization of BPE tokenizer, which leads to entirely dissimilar token representations; (b) is the Mut-BPE tokenizer introduction and our experiments framework

### C. Mutation-Aware BPE Tokenization

In variant effect prediction (VEP), the model typically compares a reference sequence and an alternative sequence that differ by a single-nucleotide variant (SNV). When standard BPE tokenization is applied independently to each sequence, even a single-base change can result in entirely different tokenizations, making it difficult for the model to isolate the effect of the mutation.

Mut-BPE addresses this issue by introducing a localized modification to the tokenization process near the mutation site (Figure 2b). It ensures that the reference and alternative sequences are tokenized identically except at the mutated position, enabling more consistent and interpretable representations for SNV comparison.

Given a pair of sequences—a reference and an alternative—that differ at a single nucleotide, we first tokenize the reference sequence using the model’s original BPE tokenizer. We then identify the token that contains the mutation site, referred to as the mutation token. This token is split into three parts: (1) the prefix (the substring before the variant), (2) the single base at the mutation site, and (3) the suffix (the substring after the variant).

The same split is then applied to the corresponding region in the alternative sequence, with the central base replaced by the alternative allele. All other tokens remain unchanged in both sequences.

For example, consider the reference sequence ATCAATGGCAATT and the alternative sequence ATCAATAGCAATT, where the base `G` at position 7 is mutated to `A`. Suppose the BPE tokenizer segments the reference as [‘ATCA’, ‘ATGGC’, ‘AATT’], with the mutation occurring within `‘ATGGC’`. Mut-BPE splits this token into `‘AT’, ‘G’`, and `‘GC’`, resulting in:

`Ref: [‘ATCA’, ‘AT’, ‘G’, ‘GC’, ‘AATT’]`

`Alt: [‘ATCA’, ‘AT’, ‘A’, ‘GC’, ‘AATT’]`

This transformation guarantees that both sequences share the same token structure except at the mutated base. The resulting tokenized sequences are then passed into the foundation model for embedding extraction and downstream VEP. In this work, we apply Mut-BPE to DNABERT-2, though the method is generally compatible with any BPE-based genomic model.

### D. Evaluation Metrics

To assess model performance in variant effect prediction (VEP), we report both the Area Under the Receiver Operating Characteristic Curve (AUROC) and the Area Under the Precision-Recall Curve (AUPRC). These complementary metrics provide a comprehensive evaluation of a model’s discriminative ability in binary classification settings.

AUROC measures the trade-off between true positive rate and false positive rate across decision thresholds, offering an overall view of ranking performance. AUPRC, by contrast, evaluates precision versus recall and is particularly informative in imbalanced classification tasks, where positive examples are rare. In such cases, AUROC can appear deceptively high, while AUPRC better reflects a model’s ability to correctly retrieve minority class instances with minimal false positives.

Several of our benchmark datasets exhibit substantial class imbalance—including the COSMIC Frequent vs. gnomAD Missense Variant dataset (183 positives vs. 15,400 negatives), the Mendelian Traits dataset (approximately 1:9 ratio), and the Complex Traits dataset (approximately 1:9 ratio). Therefore, we report both AUROC and AUPRC throughout, with particular emphasis on AUPRC for evaluating performance in imbalanced scenarios.

### E. Zero-shot based Variant Effect Prediction

In the zero-shot VEP experiments, we evaluate a model’s intrinsic ability to predict variant pathogenicity or impact without explicit fine-tuning on a task-specific training set. This is achieved by directly assessing the spatial relationships between embedding vectors generated by the models for reference and alternate DNA sequences within each dataset’s test set. The predictive performance in this setting is then quantified by calculating AUROC and AUPRC values based on these computed spatial distances, which serve as the model’s “zero-shot score.”

We compute zero-shot values for both single-DNA-strand and double-DNA-strand scenarios across all models and methods, using a sequence length of 1536 bp centered at the mutation site. For the single-strand scenario, we obtain the embedding vectors for both the reference and alternative sequences. These embeddings are then averaged along the sequence length dimension, yielding a single vector (the average token embedding) with the dimension of the model’s hidden layer. Spatial distances are subsequently computed between these averaged vectors. For the double-strand scenario, we independently calculate embeddings for the forward and reverse-complement strands. These two embeddings are then concatenated to form a single vector, after which spatial distances are computed. AUROC and AUPRC values derived from these spatial distances are then reported as the model’s zero-shot score for the test set, reflecting its performance without exposure to task-specific training data.

To provide a multifaceted assessment of the embedding space, we employ several distance metrics to quantify the relationship between the embedding vectors of the reference and alternate sequences: Cosine Similarity, which measures angular similarity indicating directional alignment in high-dimensional spaces; Pearson Similarity (Correlation Coefficient), which quantifies the linear correlation reflecting proportional changes; and Spearman Similarity (Rank Correlation Coefficient), which assesses the monotonic relationship (rank correlation) providing robustness to outliers and non-linear transformations.

### F. Fine-tune based Variant Effect Prediction

In the fine-tuning based VEP experiments, we employ full-parameter fine-tuning for all selected models with the one hidden-layer binary-classification head. This strategy is chosen given their typical parameter counts (around 100 million), allowing for maximal flexibility and comprehensive task-specific adaptation by updating all model weights during training.

For most fine-tuning experiments, the input sequence length is set to 768 base pairs (bp), centered at the mutation site, for each individual sequence in both single-strand and doublestrand scenarios. For training, we concatenate the reference sequence and the alternative sequence along the sequence length dimension. This results in an effective training sequence length of 1536 bp when only forward strands are considered. When reverse-complement strands are enabled, the training sequence length extends to 3072 bp due to the concatenation of both forward and reverse-complement pairs. To mitigate computational resource limitations and out-of-memory issues associated with longer sequences and larger datasets, a specific adjustment is made for the double-strand fine-tuning experiments on the causal eQTL dataset: for this dataset, the length of each individual sequence is reduced to 384 bp, resulting in a total training sequence length of 1536 bp for the double-strand configuration.

In fine-tuning experiments, we use a learning rate of 1 × 10^−4^ and train for 3 epochs across all settings, except for the double-strand causal eQTL dataset, which uses a reduced learning rate of 3 × 10^−5^. We apply a dropout rate of 0.1, a batch size of 16, and use the AdamW optimizer for all experiments.

### G. Foundation Models for Variant Effect Prediction

To evaluate the effectiveness of Mut-BPE in variant effect prediction (VEP), we compare it against six genomic foundation models that employ diverse tokenization strategies and architectural designs. These models serve as benchmarks across all experimental settings and datasets.

As our Mut-BPE model is based on DNABERT-2, we set DNABERT-2 [11] with its original BPE tokenizer as our primary baseline model, which allows us to directly assess the effect of substituting Mut-BPE for standard BPE within the same model architecture.

We also include GENA-BERT-Base-LastLN-T2T from the Gena-LM family [12], a BPE-based transformer model optimized for long input sequences. Due to input length constraints, this model uses a maximum sequence length of 1536 bp in all double-strand fine-tuning experiments.

To represent k-mer-based tokenization, we evaluate NTv2-100M-Multi from the Nucleotide Transformer (NT) family [27], a foundation model pre-trained on a broad set of human and multi-species genomes. Its use of fixed-length k-mers enables robust performance across a variety of downstream genomic tasks.

We additionally consider three character-level models. Hyena-Medium-450k-seqlen, from the HyenaDNA family [26], is based on an implicit convolution architecture. Caduceus-Ph [25], from the Caduceus family, is a bidirectional, reverse-complement-equivariant model built on the Mamba architecture. MutBERT [28], a probabilistic genome-based masked language model, is specifically designed to capture SNP effects from population-scale variation data.

All models are evaluated under the same experimental framework, which includes zero-shot inference and fine-tuning. This comprehensive comparison enables us to isolate the effect of tokenization strategy and assess the benefits of Mut-BPE relative to other model families.

## III. Experiments and Results

### A. Zero-shot Results

We conducted zero-shot variant effect prediction (VEP) experiments across multiple genomic foundation models, evaluating their performance using cosine similarity, Pearson correlation, and Spearman correlation metrics. Each model was assessed under both the single-strand (forward strand only) and double-strand (including reverse complement) settings. For balanced datasets, performance is reported using AUROC, while for imbalanced datasets, AUPRC is used. Detailed results are presented in Table I and Table II, respectively.

**TABLE I.** Zero-SHOT Average Results for Balanced Datasets (AUROC)

| Model | Cosine |  | Pearson |  | Spearman |  |
| --- | --- | --- | --- | --- | --- | --- |
|  | Single | Double | Single | Double | Single | Double |
| NTv2-100M-Multi | 0.445 | 0.445 | 0.452 | 0.445 | 0.448 | 0.441 |
| Caduceus-Ph | 0.512 | 0.512 | 0.500 | **0.516** | 0.489 | 0.494 |
| HyenaDNA | 0.464 | 0.464 | 0.466 | 0.464 | 0.466 | 0.464 |
| Gena-LM | 0.493 | 0.493 | **0.511** | 0.506 | **0.513** | 0.506 |
| DNABERT-2 | 0.459 | 0.459 | 0.469 | 0.459 | 0.459 | 0.458 |
| Mut-BPE | **0.514** | **0.514** | **0.511** | 0.514 | 0.507 | **0.515** |

**TABLE II.** Zero-SHOT Average Results for Imbalanced Datasets (AUPRC)

| Model | Cosine |  | Pearson |  | Spearman |  |
| --- | --- | --- | --- | --- | --- | --- |
|  | Single | Double | Single | Double | Single | Double |
| NTv2-100M-Multi | 0.065 | 0.065 | 0.066 | 0.065 | 0.068 | 0.066 |
| Caduceus-Ph | 0.070 | **0.083** | 0.080 | **0.085** | 0.062 | 0.066 |
| HyenaDNA | 0.066 | 0.063 | 0.065 | 0.064 | 0.066 | 0.064 |
| Gena-LM | 0.068 | 0.075 | 0.067 | 0.075 | 0.066 | 0.073 |
| DNABERT-2 | 0.074 | 0.072 | 0.070 | 0.074 | 0.069 | 0.075 |
| Mut-BPE | **0.088** | 0.079 | **0.087** | 0.079 | **0.085** | **0.077** |

The Mut-BPE method consistently demonstrates strong performance across both balanced and imbalanced datasets. Given the zero-shot setting, in which models are evaluated without any task-specific fine-tuning, Table I shows that most models—including Mut-BPE—achieve AUROC scores near 0.5 on balanced datasets. Table II reports performance on severely imbalanced datasets, where all models achieve AUPRC scores below 0.1. Mut-BPE outperforms other models and methods in 8 of the 12 totall evaluated scenarios. In the remaining 4 cases, Mut-BPE consistently ranks as the second-best performing model within that specific scenario (Table I, II), underscoring its robust generalizability. Despite this challenging scenario, Mut-BPE consistently outperforms other models, indicating that its embeddings more effectively capture the impact of single-nucleotide variants even without training.

Visualizing these trends in Figure 3, which showcases the performance on causal eQTL and Mendelian traits datasets, further highlights Mut-BPE’s superiority. Our method emerges as the best-performing approach in 8 of 12 the displayed scenarios. Mut-BPE exhibits a comprehensive advantage in the double-strand setting for the causal eQTL dataset and in single-strand setting for the Mendelian traits dataset. Notebly, when using Spearman as the distance metric, Mut-BPE achieves an AUPRC of 0.17 in the Mendelian traits double-strand experiments, surpassing the second-best DNABERT-2 model’s AUPRC of 0.116 by over 45%.

**Fig. 3.**
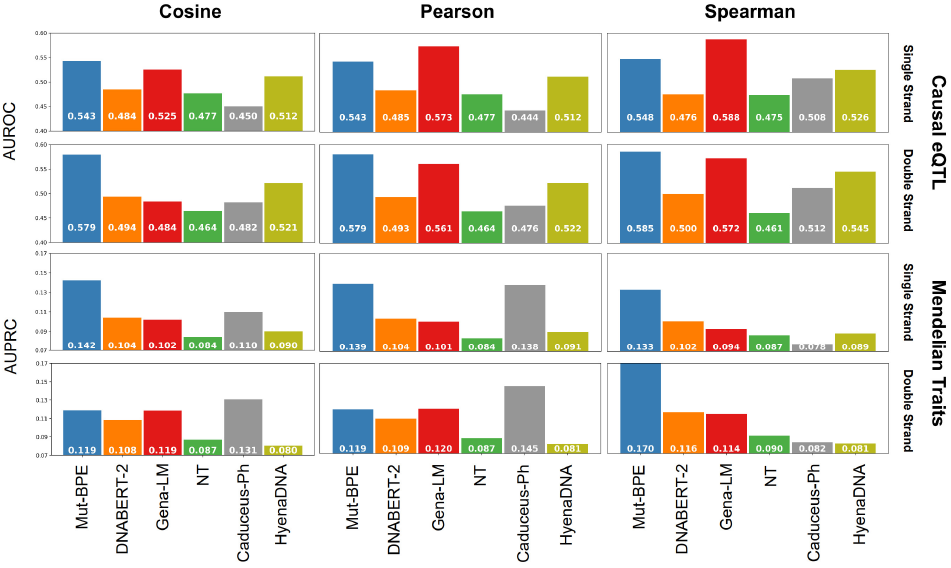
Zero-shot performance of models on the causal eQTL and Mendelian traits datasets in both single-strand and double-strand scenarios

These results suggest that equipping BPE-based models with single-nucleotide resolution via Mut-BPE improves their ability to encode mutation-induced changes at the embedding level. Compared to DNABERT-2 with its original BPE tokenizer, Mut-BPE achieves an improvement of 11.29% in average AUROC on balanced datasets and an average AUPRC improvement of 14.06% on imbalanced datasets.

### B. Fine-tuning Results

We conducted fine-tuning experiments on Mut-BPE and six other pre-trained genomic large language models with different tokenizers, across six distinct datasets, in both single-strand and double-strand scenarios. As illustrated in Figure 4, which presents fine-tuning results for three representative datasets (ClinVar Pathogenic, COSMIC Frequent vs. gnomAD Missense, and Mendelian Traits) in single-or double-strand configuration, our proposed training-free Mut-BPE method consistently performs well. Mut-BPE ranks within the top two models across all displayed scenarios. Notably, it achieves superior performance across all AUPRC metrics, once again demonstrating its strong capability in identifying significant mutations that can cause substantial human impact.

**Fig. 4.**
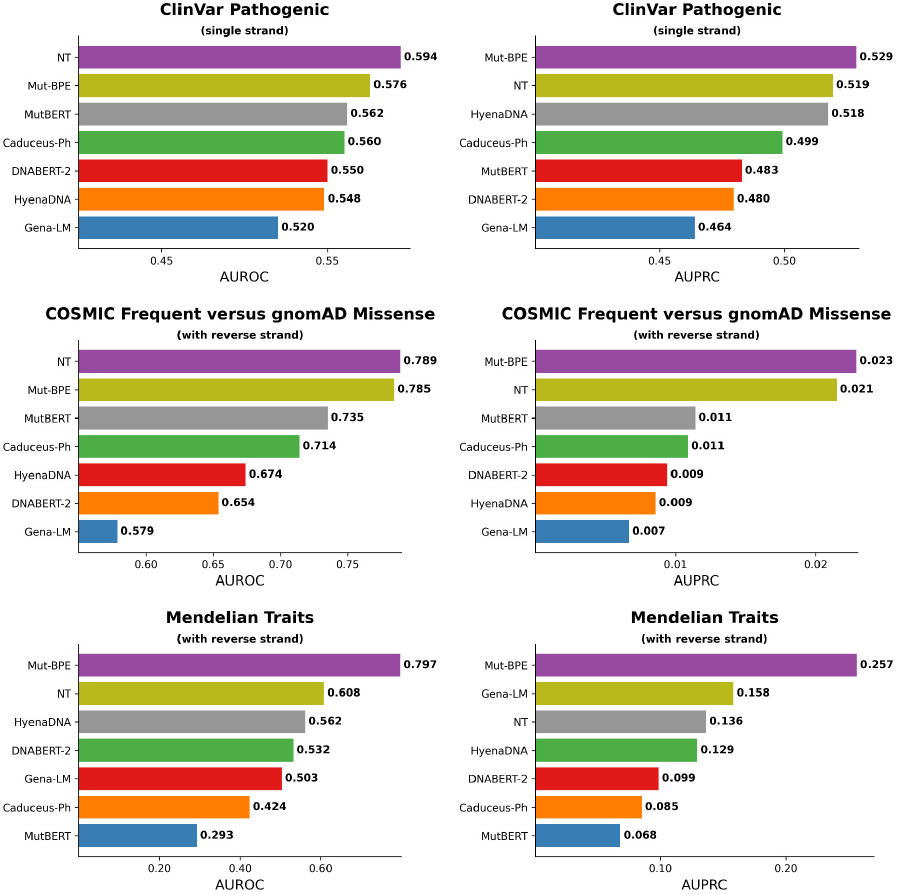
Fine-tuning results of three representative datasets (ClinVar Pathogenic, COSMIC Frequent vs. gnomAD Missense, and Mendelian Traits) in single- or double-strand configuration

We also report the overall average AUROC and AUPRC for these models and methods across all six datasets, as shown in Table III. Despite being a training-free method built upon DNABERT-2, Mut-BPE consistently achieves the best performance across all fine-tuning metrics. In single-strand experiments, Mut-BPE reaches an AUROC of 0.6418, surpassing the second-best MutBERT (which is specifically trained for single nucleotide polymorphisms) by a notable margin. Its AUPRC in single-strand also reaches 0.3576, out-performing the NT model, which has a comparable parameter count. In double-strand experiments, our method maintains its leading edge, even surpassing the excellent Caduceus model, which simultaneously supports bi-directionality and reverse-complement equivariance. Mut-BPE’s AUROC and AUPRC in double-strand are 0.6712 and 0.3739 respectively, outperforming the second-best NT.

**TABLE III.** Fine-TUNE Average Results Summary.

| Model | Single Strand |  | Double Strands |  |
| --- | --- | --- | --- | --- |
|  | AUROC | AUPRC | AUROC | AUPRC |
| NTv2-100M-Multi | 0.6237 | 0.3495 | 0.6579 | 0.3639 |
| Caduceus-Ph | 0.6098 | 0.3406 | 0.6225 | 0.3492 |
| HyenaDNA | 0.6117 | 0.3419 | 0.6112 | 0.3482 |
| MutBERT | 0.6344 | 0.3391 | 0.6357 | 0.3413 |
| Gena-LM | 0.5776 | 0.3039 | 0.5479 | 0.3097 |
| DNABERT-2 | 0.6055 | 0.3224 | 0.5943 | 0.3333 |
| Mut-BPE | **0.6418** | **0.3576** | **0.6712** | **0.3739** |

Analyzing the performance of DNABERT-2 and Gena-LM, which utilize traditional BPE tokenizers, reveals relatively lower VEP performance, often with AUROC values not exceeding 60%. This suggests that models without specific resolution for single-nucleotide polymorphic mutations may struggle in VEP tasks. However, when comparing our Mut-BPE method to the traditional BPE-tokenized DNABERT-2 model, Mut-BPE shows significant improvements across all four metrics: an approximate average increase of 6% in single-strand AUROC, 10.92% in single-strand AUPRC, 12.93% in double-strand AUROC, and 12.18% in double-strand AUPRC. This demonstrates that even without additional training and using identical training parameters, merely by improving the tokenizer to provide the model with the objective conditions to learn single-nucleotide mutation patterns, substantial progress can be achieved in VEP tasks. Overall, Mut-BPE’s performance in this experiment is excellent, with ranking first place among all methods. Its outstanding AUPRC performance in imbalanced datasets further highlights Mut-BPE’s superior accuracy in predicting positive classes, indicating its ability to more precisely identify deleterious mutations that cause disease or significantly impact phenotypes compared to other tokenizers and models.

## IV. Disscussion

In this work, we presented Mut-BPE, a simple yet effective modification to BPE tokenization that enables single-nucleotide resolution in variant effect prediction (VEP) tasks. This method is training-free, model-agnostic, and easily integrates with any BPE-based genomic foundation model. Using DNABERT-2 as a representative backbone, we evaluated Mut-BPE across six diverse mutation datasets under two experimental settings: zero-shot inference and fine-tuning. In all settings, Mut-BPE consistently outperformed standard BPE tokenization, demonstrating improved ability to capture the effects of single-nucleotide variants at the embedding level and enhancing overall VEP performance.

While these results highlight the robustness and general utility of Mut-BPE, our current evaluation is limited to a single model backbone. Future work will explore the generalizability of Mut-BPE across a broader range of BPE-based genomic models. In addition, we aim to investigate the underlying mechanisms by which Mut-BPE improves variant representations, through embedding visualization, attention pattern analysis, and gradient-based interpretability techniques. Such insights may inform the design of even more effective tokenization strategies tailored to mutation-aware genomic modeling.

## Acknowledgment

Anonymous.

